# Genome-wide SNPs of vegetable leafminer, *Liriomyza sativae*: insights into the recent Australian invasion

**DOI:** 10.1101/2022.01.06.475194

**Authors:** Xuefen Xu, Tom L. Schmidt, Jiaxin Liang, Peter M. Ridland, Jessica Chung, Qiong Yang, Moshe Jasper, Paul A. Umina, Ary A. Hoffmann

## Abstract

*Liriomyza sativae*, the vegetable leafminer, is a serious agricultural pest originally from the Americas which has now colonized all continents except Antarctica. In 2015, *L. sativae* arrived on the Australian mainland and established on the Cape York Peninsula in the northeast of the country. Here, we assessed genetic variation in *L. sativae* based on genome-wide single nucleotide polymorphisms (SNPs) generated by double-digest restriction-site associated DNA sequencing (ddRAD-seq) to uncover the potential origin(s) of this pest in Australia and contribute to reconstructing its global invasion history. Our principal component analyses (PCA) results suggested that Australian mainland populations were genetically close to populations from the Torres Strait and had connections to Bali and Papua New Guinea (PNG), whereas populations from Asia and Africa were more distantly related. Hawaii was genetically distinct from populations from Asia, Africa and Australia. Co-ancestry analyses pointed to signals of gene flow from the Torres Strait into the Australian mainland, while Indonesia/PNG were the likely sources of the initial invasion into the Torres Strait. Admixture analyses further revealed that *L. sativae* from the Torres Strait had genetic diversity originating from multiple sources, which has now spread to the Australian mainland. The *L. sativae* lineages from Asia/Africa appear closely related and may share co-ancestry. Isolation by distance (IBD) was found at a broad global scale, but not within small regions, suggesting human-mediated factors contribute to the local spread of this pest. Overall, our findings highlight the challenges in quarantine measures aimed at restricting the distribution of this global pest.

## 1 INTRODUCTION

Over the past 40 years, several polyphagous *Liriomyza* (Diptera: Agromyzidae) leaf-mining species have become recognized as pests capable of causing frequent and severe outbreaks (Murphy & LaSalle, 1999). *Liriomyza sativae* Blanchard is one of these potentially damaging leaf-mining pests, with damage resulting from tunneling activities of larvae that in turn decreases the vigor and the photosynthetic capacity of host plants (Johnson et al., 1983; Parrella, 1987). Leaf punctures caused by female flies of this species also facilitate the transfer of plant disease, with studies demonstrating that female *Liriomyza sativae* can transmit viruses of the potyvirus group (Zitter & Tsai, 1977). *Liriomyza sativae* is highly polyphagous across plant families including Asteraceae, Cucurbitaceae, Fabaceae, Solanaceae, and Umbelliferae (Spencer, 1973), with its host range potentially expanding as it colonizes new areas, as in the case of an expansion to green onions in Hawaii (Carolina et al., 1992).

Although *L. sativae* originated in the Americas, it has now expanded its geographic range and colonized many areas throughout the globe (Scheffer & Lewis, 2005). The movement of infested plant material through international trade and transportation has likely facilitated this colonization process (Minkenberg, 1988). Eggs and larvae of *L. sativae* are embedded internally within plant leaves and can be easily transported from production areas to market without being noticed. The pupae of *L. sativae* may also be transported through infested soil or plant debris, while strong winds may facilitate long-distance adult dispersal (Fenoglio et al., 2018). *Liriomyza sativae* are typical secondary pests with outbreaks occurring after the indiscriminate use of broad-spectrum insecticides, which removes natural enemies as well as promoting the evolution of resistance (Mason et al., 1987; Oatman & Kennedy, 1976; Parrella & Keil, 1984; Reitz et al., 2013; Ridland et al., 2020). Once *L. sativae* invade new regions, it is difficult to eradicate it (Scheffer & Lewis, 2005), highlighting the importance of quarantine.

Following invasion and establishment into countries, *L. sativae* has often had an immediate and detrimental impact on local horticultural industries (Murphy & LaSalle, 1999). For example, the *Liriomyza sativae* invasion led to substantial economic damage in China where the pest was first found in 1993 and then expanded into most agricultural areas, with over 2.7 million hectares affected by the end of 2005 responsible for a loss of around 3 billion yuan annually (Chunlin et al., 2005). *Liriomyza sativae* was first detected in Australia in 2008 when established populations were identified in the Torres Strait (Blacket et al., 2015). Subsequently, *L. sativae* was recorded at Seisia on the northern tip of the Cape York Peninsula, located in the northeastern part of the Australian mainland in 2015 (IPPC, 2017). Modeling suggests that further spread of this pest is likely on the mainland unless it is successfully prevented by quarantine (Maino et al., 2019). Pest activity is expected to overlap with the production cycle in Australia of several high-risk horticultural crops in the vegetable, production nursery, and melon industries (Maino et al., 2019).

Understanding the invasion history of pests like *L. sativae* can help inform quarantine and management actions, and predict the likelihood of insecticide resistance genes entering colonizing populations (Ma et al., 2007). The invasion history of a pest can be determined not only from historical records but also through genetic approaches. To date, information on the genetic structure of *L. sativae* populations is limited. Previous research on genetic markers in this species mainly focused on mitochondrial DNA (mtDNA) for deciphering *L. sativae* population structure (Blacket et al., 2015; Parish et al., 2017; Scheffer & Lewis, 2005; Tang et al., 2016; Xu et al., 2021a). While mtDNA markers provide some indication of the relatedness of populations, they have a relatively low resolution particularly in detecting fine-scale structure (Anderson et al., 2010). Patterns of population relatedness based on mtDNA can also be obfuscated by processes like endosymbiont invasions and recombination across the mitochondrial genomes (Ballard & Whitlock, 2004).

With the increasing ease and speed of DNA sequencing, high-resolution genomic SNP-based methods like double digest restriction site-associated DNA sequencing (ddRAD-seq) provide a new dimension to population-level studies of species (Peterson et al., 2012). These markers have been successfully applied to understand patterns of movement in insects that act as pests or disease vectors (e. g. Ryan et al., 2019; Schmidt et al., 2019; Yan et al., 2021). They provide a much higher level of resolution than other nuclear marker systems like microsatellites given the sheer number of marker loci that can be scored (Fola et al., 2020). SNP markers scattered across the genome also provide the opportunity to use genetic approaches in tackling new questions such as estimating cross-generation movement rates of pests following SNP-based identification of related individuals (Cordeiro et al., 2019; Jasper et al., 2021), and the identification of genomic regions that may be under strong selection such as those linked to pesticide resistance (Endersby-Harshman et al., 2020; Uchibori-Asano et al., 2019; Yang et al., 2020).

In this study, we use SNP-based methods to investigate in detail the incursion patterns of *L. sativae* into the Cape York Peninsula region in Australia. A large panel of SNPs were generated by ddRAD-seq and screened across 24 *L. sativae* populations, focusing mostly on the Asia-Pacific region but also including some worldwide samples. This study aimed to: (1) explore the population genetic structure of *L. sativae* across this region, (2) determine the possible source(s) of the recent incursions in Australia, (3) estimate genetic diversity across different *L sativae* populations, including older and more recent invasions, and (4) provide a baseline for future explorations of further *L. sativae* incursions across the mainland.

## 2 MATERIALS AND METHODS

### 2.1 Sample collections, DNA extraction, DNA barcode

*Liriomyza sativae* field samples were obtained from 24 populations from 8 countries between July 2016 and May 2020 (Fig. 1, Table 1). Samples were preserved in 100% ethanol at −80°C until DNA extraction. As species of *Liriomyza* have similar external morphology, taxonomic identification of *Liriomyza sativae* samples was confirmed by both morphological identification (by Dr. Mallik Malipatil, Agriculture Victoria, AgriBio, La Trobe University, Bundoora, Australia) and barcodes based on mitochondrial CO1. For the barcoding, legs of flies were taken from specimens for Chelex DNA extraction with protocols following previous work except in 70 μL of 5% Chelex 100 resin (Bio-Rad Laboratories) (Coquilleau et al., 2021; Xu et al., 2021a,b). Once species identity was confirmed, whole fly bodies were used for ddRAD-seq libraries.

**FIGURE 1.**
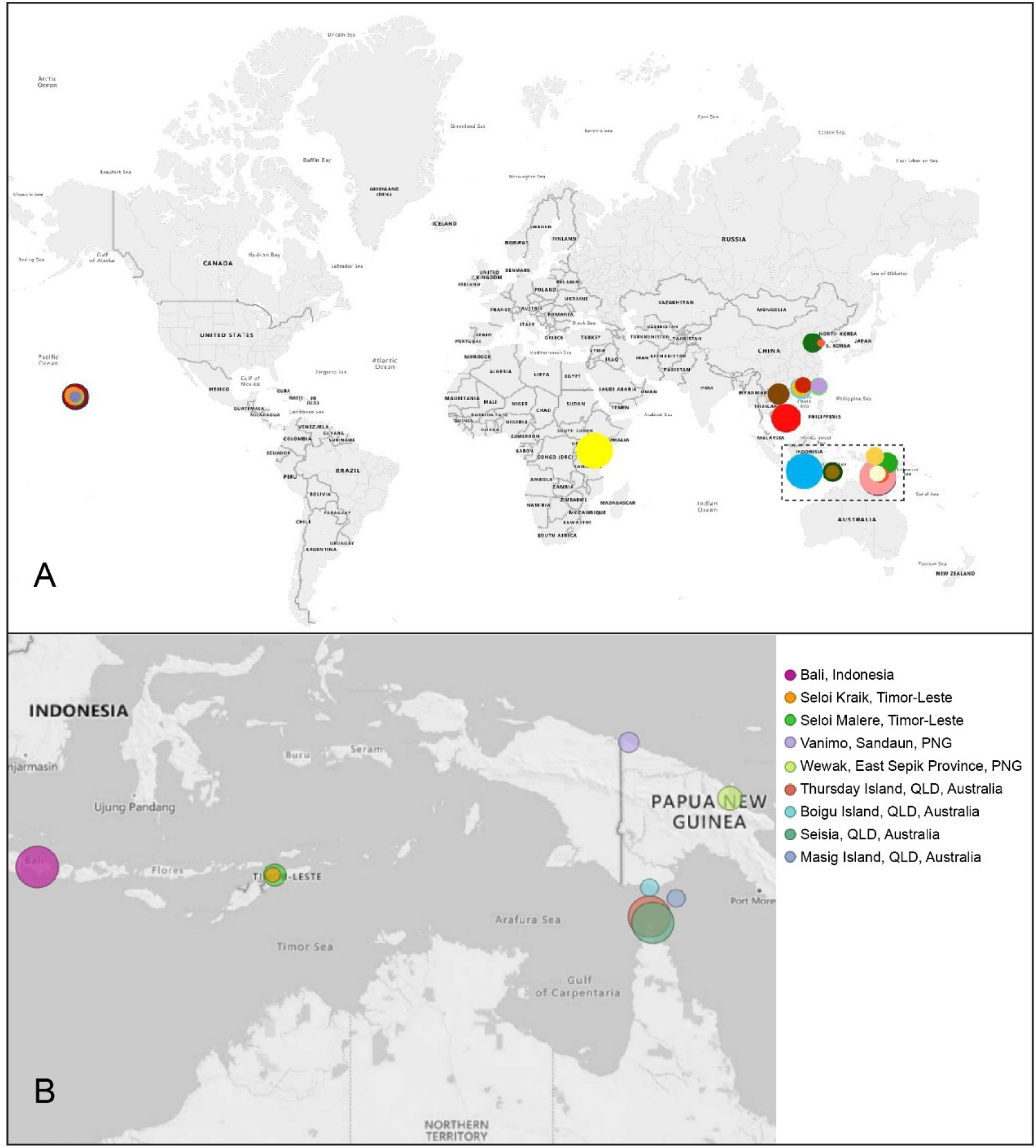
(A) Sampling locations of *Liriomyza sativae* in this study. Colours indicate different populations and the size of circles reflect the relative sizes of samples. The dotted rectangle represents the local geography around northern Australia, as detailed in (B).

**Table 1.**
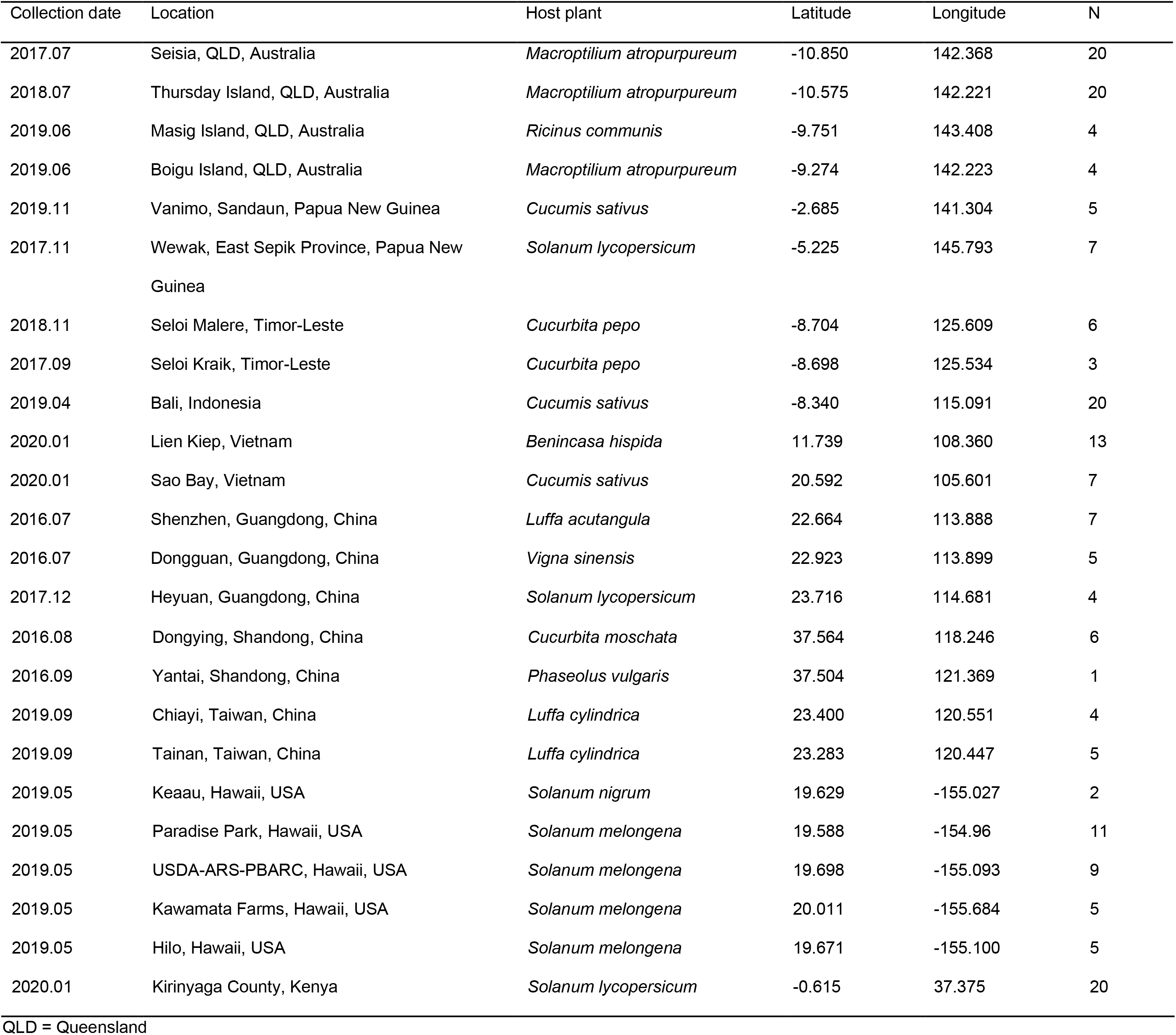
The 193 *Liriomyza sativae* samples retained after quality filtering for ddRAD sequencing.

### 2.2 ddRAD-seq libraries preparation

Genomic DNA from individual *L. sativae* was extracted with a DNeasy Blood and Tissue kit (Qiagen, Venlo, Limburg, NL). ddRAD-seq libraries were prepared according to the protocol from Rašić et al., (2014). In brief, around 45ng of high-quality genomic DNA was simultaneously doubled-digested using restriction enzymes *MluC*I (10 units, New England Biolabs, Beverly MA, USA) and *Nla*III (10 units, New England Biolabs, Beverly MA, USA), NEB CutSmart buffer, and water. Digestions were incubated at 37°C for 3 h and then the products were cleaned with 1.5× volume of AMpure XP paramagnetic beads (Beckman Coulter, Brea, CA, USA). After that, the digestion fragments were ligated to modified Illumina P1 and P2 adapters overnight at 16°C with T4 ligase (1000 units, New England Biolabs, Beverly, MA, USA), followed by a heat deactivation step at 65°C for 10 minutes. Adapter ligated DNA fragments from all individuals were pooled and cleaned with 1.5× bead solution afterward. Pippin-Prep 2% gel cassette (Sage Sciences, Beverly, MA) were applied for size selection of fragments between 350-450 bp. Lastly, 1 μL size-selected DNA was added in a 10 μL PCR reaction as a template with 5 μL of the Phusion High Fidelity 2× Master mix (New England Biolabs, Beverly MA, USA) and 2 μL of 10 μM P1 and P2 primers. PCR programs were 98°C for 30 s, 15 cycles of 98°C for 10 s, 60°C for 30 s, 72°C for 90 s, and final elongation at 72°C for 5 min. We pooled ten such PCR reactions together and cleaned them with a 1.5× bead solution for making the final library. The ddRAD-seq libraries were sequenced on the Illumina HiSeq2500 platform to obtain 150 bp paired-end reads.

### 2.3 Sequence data processing and genotyping

We processed the raw sequencing reads according to the STACKS v2.54 pipeline (Catchen et al., 2013). Quality filtering of raw reads was performed with the process_radtags program. This program first checks if the barcode and the RAD cut site are intact, and then demultiplexes the data, correcting errors within allowable limits. We first trimmed the reads to 115 bp in length, and removed sequence reads with average Phred scores below 20, a cut-off used in similar studies (Chen et al., 2021; Schmidt et al., 2019). Given the absence of a reference genome for *Liriomyza sativae*, a STACKS catalog was assembled *de novo* using the denovo_map.pl program, following the standard workflow of *de novo* assembly for diploids (Catchen et al., 2011). After catalog construction, we removed 7 samples due to a high level of missing data (>50%) and generated VCF files containing SNPs called in 75% of individuals from each population with a minor allele count of 3 for downstream analyses (c. f. Hoffmann et al., 2021).

### 2.4 POPULATION STRUCTURE

#### 2.4.1 Principal component analysis

For principal component analysis (PCA), we used the “dudi.pca” function implemented in the ade4 R package to create a genotype covariance matrix (Dray & Dufour, 2007). We ran two PCA analyses, summarizing genotypic variation across all populations and all populations except Hawaii (see below).

#### 2.4.2 FineRADstructure analyses

We performed a co-ancestry analysis on all *L. sativae* (193 individuals) with FineRADstructure, which is designed to infer recent population structure (available at <https://github.com/millanek/fineRADstructure>). The co-ancestry matrix of 193 individuals was based on a summary of nearest neighbor haplotype relationships, which were calculated from multiple SNPs per locus. The individuals were assigned to corresponding populations and a phylogenetic tree was constructed using the FineRADstructure MCMC clustering algorithm with default settings. R scripts fineradstructureplot.r and finestructurelibrary.r (available at <http://cichlid.gurdon.cam.ac.uk/fineRADstructure.html>) were used for visualization.

#### 2.4.3 Admixture analysis

Admixture v.1.3.0 (Alexander et al., 2009) was used to infer genetic structure assuming a specific number of ancestral lineages (K). This tool is based on a maximum likelihood method to estimate individual ancestries, and a cross-validation (CV) test was applied to evaluate the optimal K (Alexander and Lange, 2011). To avoid stochastic effects from a single analysis, we ran 10 iterations at each value of K. Data were visualized using ComplexHeatmap and viridis package in R (downloaded packages from https://www.bioconductor.org/).

#### 2.4.4 Isolation-by-distance

To test whether genetic differentiation patterns matched expectations from isolation by distance (IBD), we used a Mantel test to explore the correlation between genetic distances (F_ST_) and geographic distances (km) based on 999 permutations implemented in adegenet 2.1 (Jombart, 2008). To avoid sampling bias, we removed populations with fewer than 4 individuals (Yantai, Shandong, China; Keaau, Hawaii, USA; Seloi Kraik, Timor-Leste), leaving 21 populations for the overall IBD analysis. We also ran a region-specific IBD that included 8 populations (Bali, 1 Timor-Leste location, 2 PNG locations, Boigu Island, Masig Island, Thursday Island, Seisia) of particular interest to the recent *L. sativae* invasion into Australia. Genetic divergence was estimated by calculating F_ST_ values between all pairs of populations using the ‘genepop’ R package (Rousset et al., 2020). Geosphere R package v.1.5-5 was applied to calculate straight-line geographic distances between populations (Hijmans, 2017). P-values smaller than 0.05 were considered to be significant.

## 3 RESULTS

### 3.1 Principal component analysis (PCA analysis)

Individual relationships within and between populations were visualized using PCA analyses that contained all samples (193 individuals) (Fig. 2) and all samples excluding Hawaii (161 individuals) (Fig. 2). A total of 7379 SNPs loci were retained for 193 individuals from 24 populations (Table 2). When all individuals and sites were included in a PCA (Fig. 2), there was a strong separation of the Hawaii populations from the other areas, with the main axes accounting for 6.1 % and 3.7% of the variation.

**FIGURE 2.**
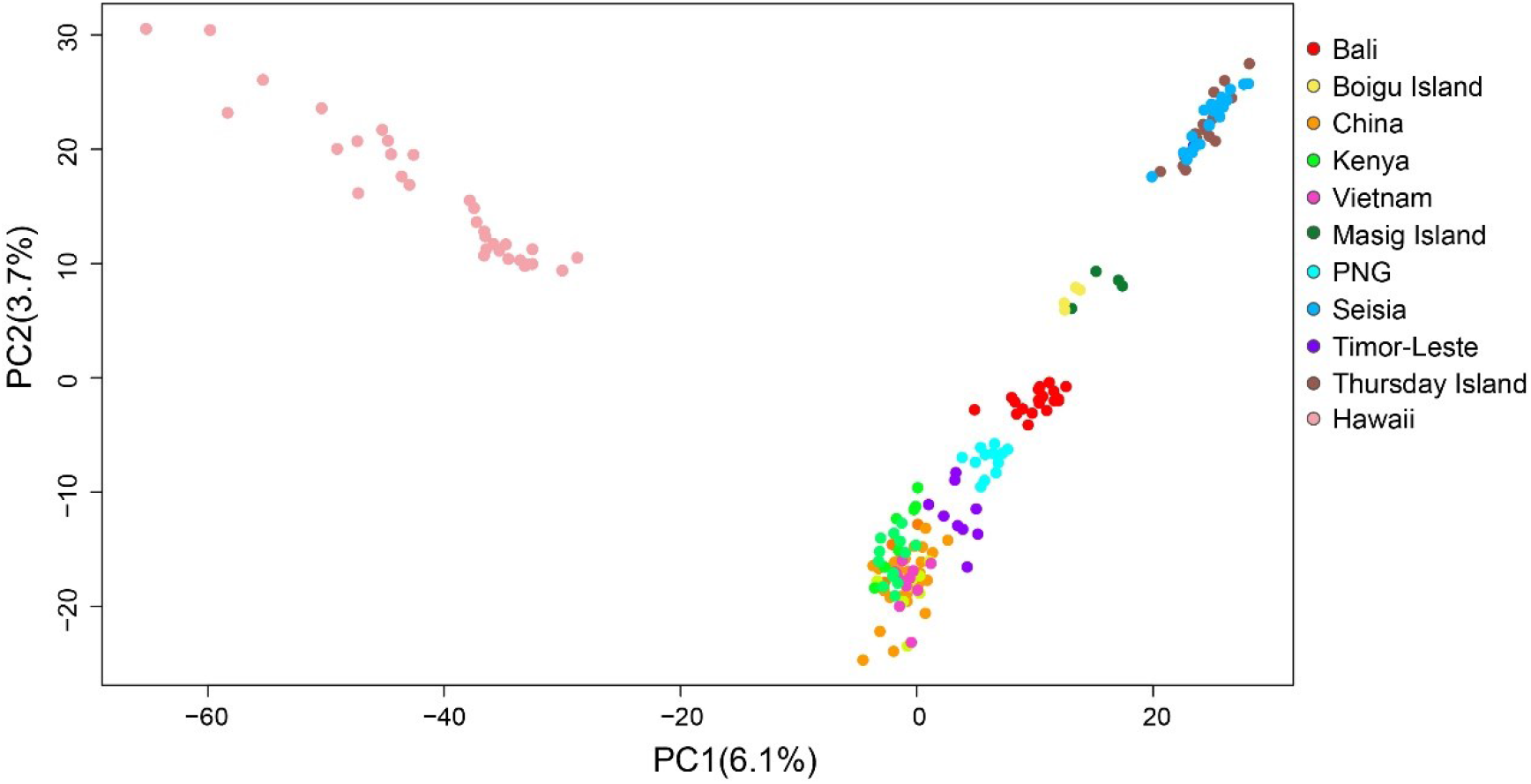
Principal components analysis (PCA) of 193 individuals collected from 24 populations based on 7,379 SNPs. The variances explained by PC1 and PC2 are given. Each dot represents sampled individuals and some populations from the same country were merged except for the Australian populations.

Given the outlier position of the Hawaii populations, we constructed a separate PCA excluding Hawaii based on 7360 SNPs for 161 individuals (Fig. 3). In this PCA the first and second principal components accounted for 5.7% and 2.3% of the observed variation, respectively. On PC1, Seisia and Thursday Island were clustered together and close to Masig Island, Boigu Island, Bali (Indonesia), Papua New Guinea, and Timor-Leste. Samples from China, Vietnam, and Kenya clustered together but away from the Australian populations. PC2 further reveals that populations from Papua New Guinea and Bali were close to two populations from the Torres Strait. These patterns indicate that *L. sativae* from mainland Australia was tightly associated with populations from the Torres Strait and may also be related to Bali and PNG. Asian and African populations of *L. sativae* were distinct from Australian populations.

**FIGURE 3.**
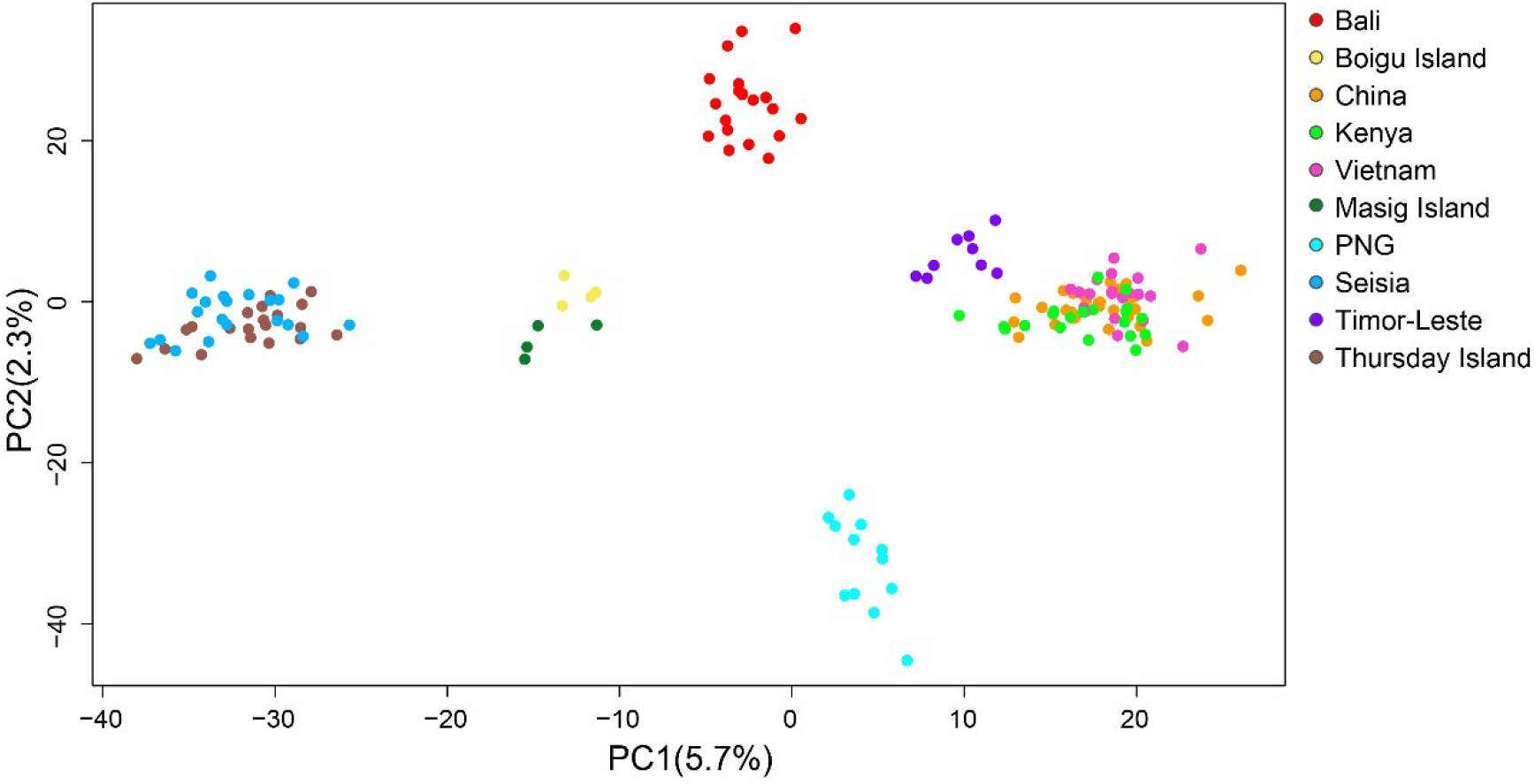
Principal components analysis (PCA) was conducted on 161 individuals based on 7,360 SNPs without the Hawaiian samples. The variances explained by PC1 and PC2 are given. Each dot represents sampled individuals and populations from the same country are combined except those from Australia.

### 3.2 FineRADstructure Analysis

The fineRADstructure plot (Fig. 4) shows four subgroups among selected locations. The biggest subgroup placed Asian genotypes together with Timor-Leste and the African location sampled, as expected from the above PCA analysis. Timor-Leste genotypes grouped with Asian countries (China and Vietnam) and clustered with Kenya, suggesting recent co-ancestry in these regions. The genotypes of Bali formed a lineage with PNG that was close to Asia/Africa clades. However, *L. sativae* from the central highland of PNG were genetically distinct from Bali, which was also confirmed by PCA analyses. The fineRADstructure analysis suggests that Bali and PNG shared recent co-ancestry but these two populations have independently genetically diverged. Genotypes from the Hawaiian populations are cluster together but with a high level of differentiation. The Hawaiian lineage was clearly differentiated from the other lineages with deep genetic divergence, indicating an invasion history of *L. sativae* in Hawaii separate from the other populations. Torres Strait populations formed a lineage with the Australian mainland population from Seisia, suggesting high rates of gene flow among these populations. Results from the FineRADstructure analysis also revealed a clear signal of invasion of the Australian mainland from the Torres Strait instead of other regions. Interestingly, Bali had the highest co-ancestry with the Torres Strait populations followed by PNG when compared to the other populations, suggesting that Bali and PNG were important sources for the initial invasion of the Torres Strait Islands.

**FIGURE 4.**
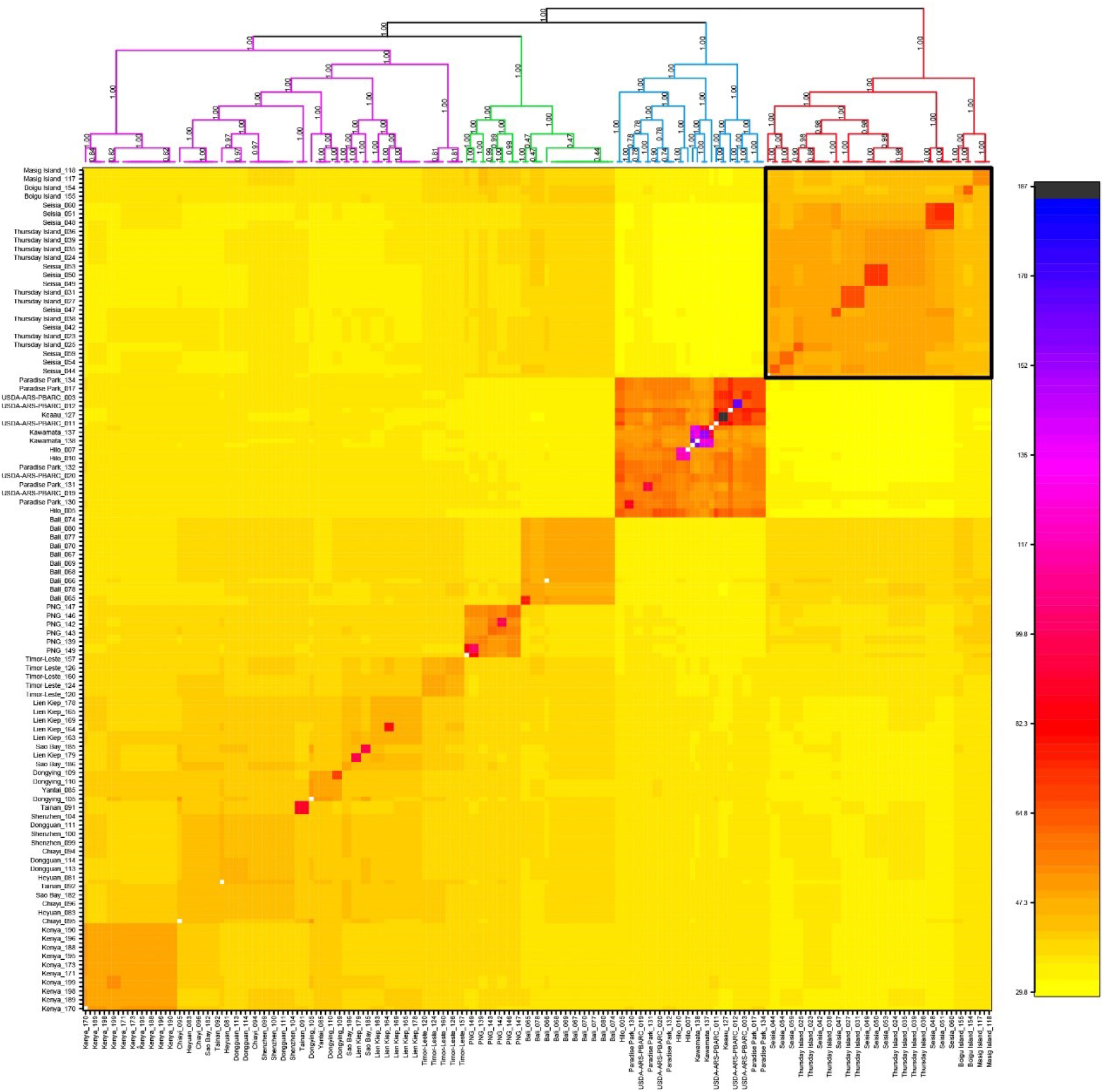
The fineRADstructure plot with co-ancestry map and phylogenetic tree. The colour scale bar indicates estimated co-ancestry, with light yellow suggesting low co-ancestry and darker yellow, orange and red indicating progressively higher co-ancestry. The Australian populations formed a clade (labelled in red) which is shown in the solid black square.

### 3.3 Admixture analyses

To infer the genetic structure and the degree of relatedness among *Liriomyza sativae* populations with different genetic backgrounds, we ran an admixture analysis with cross-validation for values of K (subgroups) from 2 through 10 to examine patterns of ancestry based on 193 individuals (Fig. S1). Note that a plot with K=1 was not considered given the low likelihood of such a scenario based on existing literature (Yamashita et al., 2019). A ten-fold cross-validation (CV) procedure was performed to infer the optimal number of subgroups. We found a minimum CV score was obtained when K=5, and we used this value in an admixture plot (Fig. 5). When K=5, individuals from Hawaii formed a distinct cluster, which was also supported in the other admixture plots (K=2-6) (Fig. S1). Individuals from Vietnam, Kenya and China shared the same ancestries and formed a cluster, which suggests these populations derived from a common origin. Timor-Leste shared the most ancestries parts from Asia/Africa and a smaller component with parts from Bali and Australian populations, The spread of *L. sativae* across Timor-Leste was likely caused by multiple introductions, thus some level of ongoing gene flow was likely during the initial stages. Unfortunately, we lack samples from southeast Asia (e.g., Malaysia, Philippines), which prevents us from fully assessing these possibilities. Further sampling in southeast countries near Timor-Leste may help reveal this part of the incursion history in more detail. Genetic variation in Hawaii, Asia and Africa was relatively low, which suggests a possible demographic bottleneck at the time of introduction and/or a period of limited population growth after establishment. The pattern of genetic differentiation across these populations indicates a limited contribution to the *L. sativae* expansion in Australia, as also highlighted by the PCA and fineRADstructure analyses. Bali and PNG are two different genetic lineages with a low proportion of putative admixture, pointing to separate introduction events. Interestingly, Bali and PNG contributed most to the ancestries of the two Torres Strait populations (Boigu Island and Masig Island), and new genotypes were detected leading to novel ancestry for Thursday Island and Seisia. Admixture results also suggest a clear signal of gene flow from the Torres Strait to mainland Australia. Compared to Seisia, *L. sativae* from the Torres Strait Islands (Boigu Island and Masig Island) contained higher levels of genetic diversity (including new genotypes) that are likely associated with multiple sources (e.g., Bali, PNG, and potentially Timor-Leste) for introductions. Our admixture results are congruent with the historic population expansion pattern of *L. sativae* in Australia, where the species was first introduced in 2008 into the Torres Strait, with a rapid expansion over multiple islands and then an introduction into the northern Cape York Peninsula area in 2015. We also show plots when K=8 (the second-best K value) indicating additional hierarchical structure levels, primarily involving Kenya, but also within Australian populations, and within Hawaii (Fig. S2).

**FIGURE 5.**
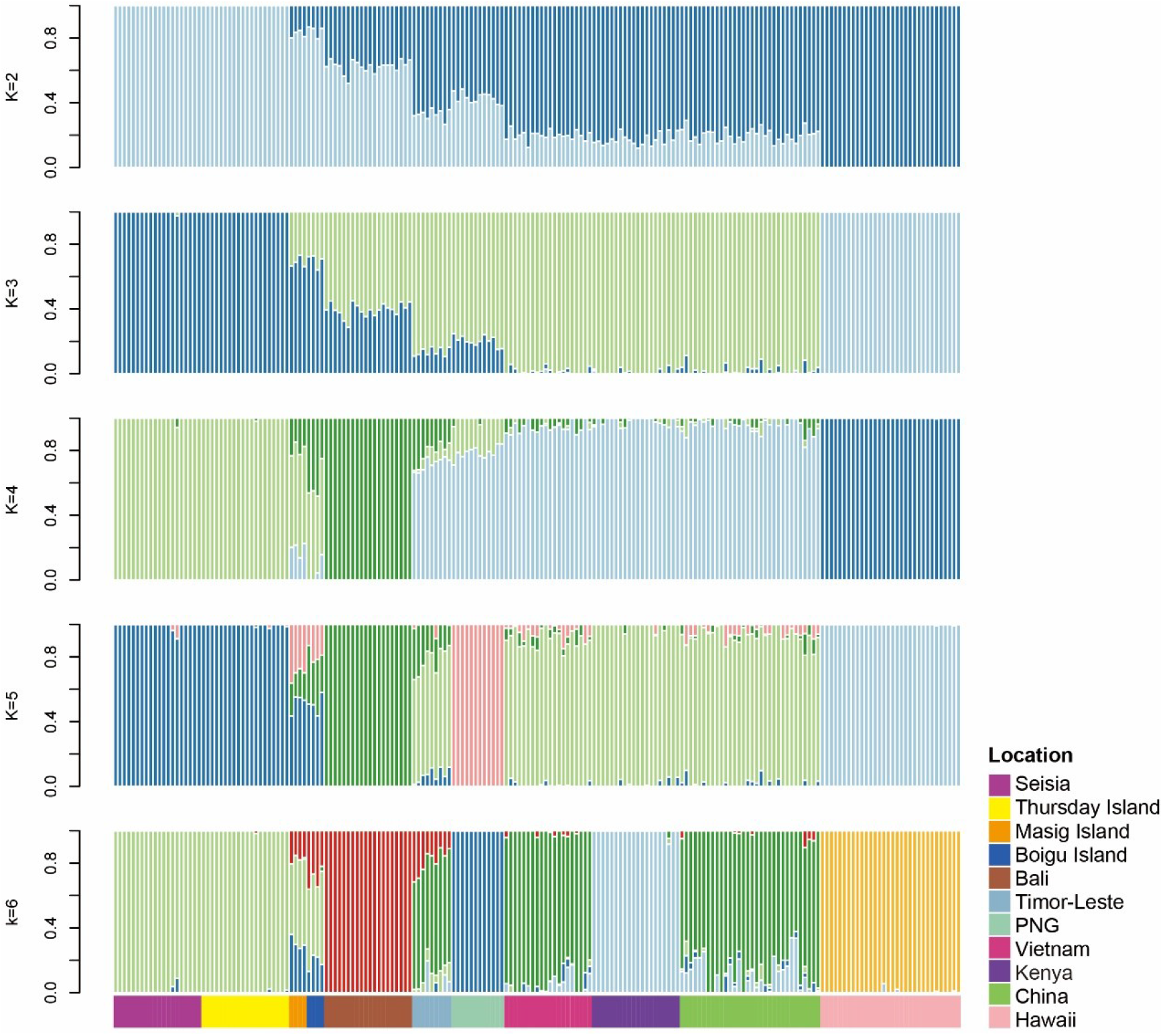
Population structure of *L. sativae* based on 193 individuals inferred using admixture analysis. The minimum K was identified as 5.

### 3.4 Isolation-by-distance

To evaluate the possible correlation between genetic and geographic distance, we built matrices of genetic distances among 21 populations (excluding 3 populations with individual numbers less than 4) by calculating pairwise F_ST_ values and then correlated these with pairwise geographic distances between populations based on 3318 SNPs. Pairwise F_ST_ values among all populations ranged from 0.0045 to 0.3269 (Table S1). Relatively stronger differentiation was observed between Kawamata Farms (Hawaii, USA) and other locations (F_ST_ values ranging from 0.148 to 0.3269). The highest F_ST_ value (0.3269) was between Kawamata Farms (Hawaii, USA) and Seisia (QLD, Australia). On the other hand, the lowest F_ST_ value (0.0045) was between Heyuan (Guangdong, China) and Chiayi (Taiwan, China). Pairwise F_ST_ values within Hawaiian regions ranged from 0.018 to 0.177, suggesting high genetic diversity within Hawaii. Specifically, the pairwise F_ST_ between Paradise Park (Hawaii, USA), Hilo (Hawaii, USA), and USDA-ARS-PBARC (Hawaii, USA) ranged from 0.0181 to 0.0502, while the pairwise F_ST_ between Kawamata Farms (Hawaii, USA) and the rest of the Hawaiian populations ranged from 0.148 to 0.1775. At a large spatial scale (21 populations excluding populations with individuals less than 4), we find support for isolation-by-distance with a Mantel test being significant (P = 0.002) based on 999 replicates (Fig. 6A). IBD results revealed a significant positive correlation between the geographic and genetic matrices in the large-scale comparison.

**FIGURE 6.**
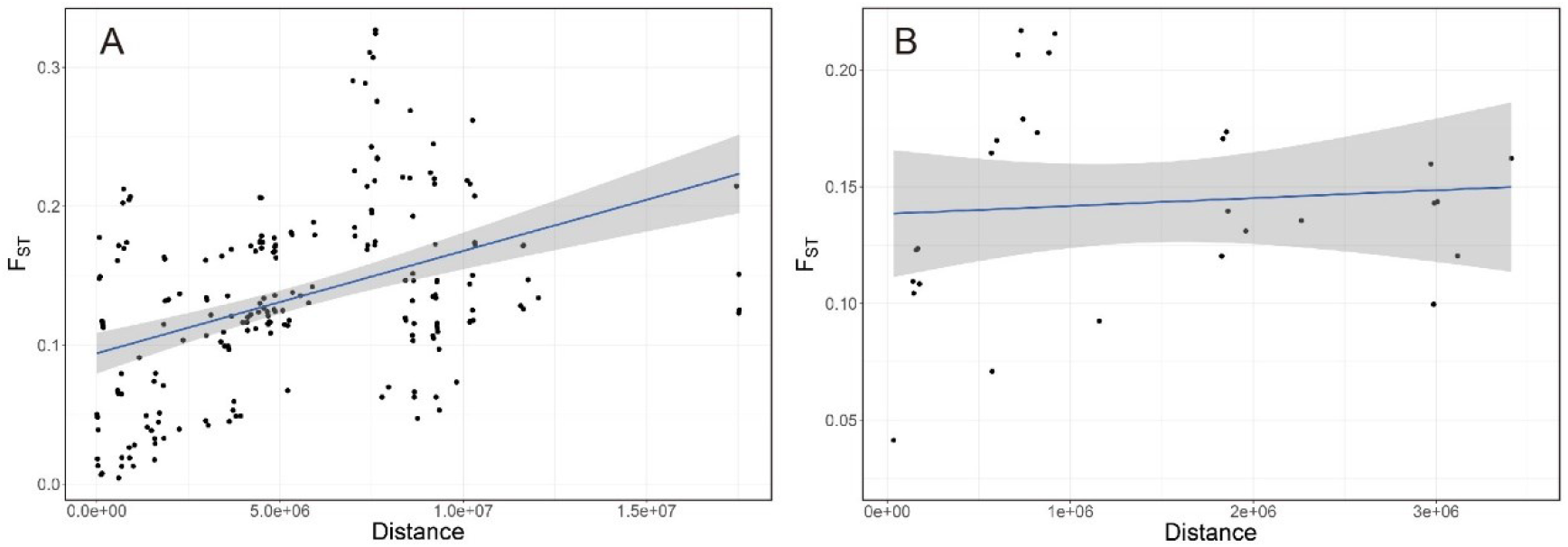
(A) Correlation analysis between genetic distances (F_ST_) and geographic distances for the 21 *Liriomyza sativae* populations; P = 0.002. (B) Correlation analysis between genetic distances (F_ST_) and geographic distances for the 8 *L. sativae* populations associated with the Australian incursion; P = 0.2169

We also examined IBD using populations that might be involved in the recent Australian *L. sativae* incursion (1 location in Indonesia, 1 location in Timor-Leste, 2 locations in PNG, and 4 locations in Australia). Pairwise F_ST_ between these populations ranged from 0.0414 to 0.2169 (Table S2). Relatively stronger differentiation occurred between two populations in PNG and the remaining populations (F_ST_ values ranging from 0.1355 to 0.2169). The highest F_ST_ value (0.2169) was between Wewak (East Sepik Province, PNG) and Seisia (QLD, Australia). On the other hand, the lowest F_ST_ value (0.0414) was between Thursday Island (QLD, Australia) and Seisia (QLD, Australia). A Mantel test indicated no relationship between pairwise genetic (F_ST_) and geographic distances (P = 0.241 based on 999 replicates) (Fig. 6B), suggesting that distance isolation had a low effect on genetic differentiation at this smaller spatial scale.

## 4 DISCUSSION

With the increasing globalization of trade and an increasing volume of goods being moved, threats posed by invasive pest species are increasing (Hulme et al., 2009). *Liriomyza sativae* represents one such threat, having invaded many countries around the world in the past 40 years (Murphy & LaSalle, 1999; Scheffer & Lewis, 2005). In Australia, *L. sativae* has now established in the north-eastern part of the mainland and it can potentially expand into other regions (Maino et al., 2019). Here, we conducted the first population genetic analyses of *L. sativae* individuals based on genome-wide SNPs, aiming to trace incursion pathways into the Australian mainland and around the region.

Based on our results, we find expected signals of gene flow from the Torres Strait into the mainland. The allelic composition of the Thursday Island (Torres Strait) population is nearly the same as that from Seisia (mainland Australia). However, these two populations are genetically different from Masig Island (Torres Strait) and Boigu Island (Torres Strait), although all these populations still cluster together as a group when compared to other populations. Our study therefore provides evidence that the incursion of *L. sativae* in the mainland of Australia is directly related to Torres Strait populations instead of incursions from other countries. We also found support for multiple invasions from different countries (especially Indonesia, PNG, and Timor-Leste) into the Torres Strait populations as reflected by results from the admixture analyses. Furthermore, we found that populations in Hawaii, USA are genetically distinct from other countries, whereas samples from Kenya, China, Vietnam, Timor-Leste clustered as a group related to Bali and PNG.

*Liriomyza sativae* (incorrectly named *Agromyza pusilla*) was damaging crops in mainland USA before World War 1 (Spencer, 1973). While *L. sativae* was first described from specimens collected from *Medicago sativa* in Argentina in 1937 (Blanchard, 1938), a South American origin for the species has not been validated with genomic markers. By 1980, *L. sativae* was common in southeast USA, Central America, and northern South America (Hardy & Delfinado, 1980). The earliest collection of *L. sativae* in Hawaii was in April 1921 (Table S3), where it was described as *Liriomyza canomarginis*, but was later synonymized with *L. sativae* (Spencer, 1973; Table S3). Given that Hawaii imports most of its vegetables from mainland USA (Parcon et al., 2010), these imported vegetables are the likely source of incursions; the CO1 haplotype in Hawaii belongs to the *sativae*-W clade (Xu et al., 2021a), which is also widespread in North America, having been recovered from crops in California, Florida, and Arizona (Scheffer & Lewis, 2005). In contrast, CO1 clade A is the most common in Florida and has not been recorded in Hawaii, suggesting that *L. sativae* in Hawaii has a west coast origin. The sativae-W clade is the invasive clade whereas all other clades are found in the Americas. Apart from highlighting the genetic uniqueness of the population from Hawaii, our data also indicate strong differentiation between the northern (Kawamata Farms) and eastern (remaining) groups in Hawaii. This strong differentiation may reflect geographic barriers since Mauna Kea, the highest point in the Hawaii Island, separates these populations.

The movement of *L. sativae* to Hawaii (first recorded in 1921) and then to several South Pacific islands (Waterhouse & Norris, 1987) occurred at least 20 years before this pest moved through Asia (Martinez, 1994) and Africa (Martinez & Bordat, 1996). In turn, from the mid-1970s, *Liriomyza trifolii* (Burgess) has largely displaced *L. sativae* in the southern United States (Parrella, 1982) and Hawaii (Johnson, 2005). This may explain why the Hawaiian leafminer populations are so distinct from the other populations in this study. The spread of *L. sativae* has speeded up since the 1990s when *L. sativae* moved eastward through tropical and sub-tropical areas like Africa, China, Indonesia and Vietnam (Andersen et al., 2002; Chen & Zhao, 1999; Rauf et al., 2000; Ridland et al., 2020). *L. sativae* arrived in Indonesia in the 1990s and became a major pest in lowland vegetable crops (Rauf et al., 2000). The species was then recorded in Timor-Leste in 2003 and West Papua in 2005. In 2008, *L. sativae* were first collected on Warraber Island in the Torres Strait in a heavily mined tomato plant (Blacket et al., 2015). Subsequently, it was recorded in the central highlands of Papua New Guinea in 2011 and other Torres Strait Islands (Blacket et al., 2015). In 2015, *L. sativae* was recorded on mainland Australia at Seisia, at the tip of Cape York (IPPC, 2017). In Kenya, *L. sativae* had became a common pest by the early 2000s (Gitonga et al., 2010; Musundire et al., 2011).

Our population genetic results are broadly congruent regarding the biogeographical history of *L. sativae*. The genotypes in Kenya, China, Vietnam, and Timor-Leste are genetically related, consistent with a relatively recent invasion across these regions. Genetic data also support a recent invasion of this species across 12 provinces in China with relatively low levels of genetic diversity (Tang et al., 2016). Despite these connections between distant locations, IBD was still detected overall, suggesting that geographical distance, wind currents, ocean currents, and obstructing landmasses at a broader scale may have affected genetic connectivity of the *L. sativae* populations (White et al., 2010; Broquet et al., 2006; Ben et al., 2017). On the other hand, when a smaller scale was considered (populations related to recent Australia incursion), genetic distances were not significantly correlated to geographic distances, suggesting that human-mediated movements may have contributed to local spread.

Although the SNP data provide a detailed picture of relatedness that is not available from mtDNA data, our results are nevertheless consistent with some previous CO1 results. These indicated a link between the Torres Strait Islands and the mainland incursion as well as the possibility of multiple invasions into Torres Strait populations (Blacket et al., 2015; Xu et al., 2021a). Previous CO1 studies demonstrate that S.27 is the dominant haplotype in Indonesian and Australian populations whereas one population (Erub Island) in the Torres Strait possesses haplotype S.07, the dominant haplotype found in PNG (Blacket et al., 2015; Xu et al., 2021a). The SNP data now provides a much clearer signature of admixture in the Torres Strait populations, including Thursday Island and Seisia which are only separated by 34 km. In future studies, it will be interesting to track further incursions into this area and beyond. Currently, the mainland population at Seisia is under quarantine to prevent its spread to other regions, but further spread seems inevitable eventually given that many other regions on the mainland are suitable for this species. Any further incursions into the Torres Strait may benefit this pest such as through the introduction of pesticide resistance genes. The geographical location of the Torres Strait is particularly suitable for multiple incursions as detected in species like the Asian tiger mosquito, *Aedes albopictus* (Beebe et al., 2013; Schmidt et al., 2021), island sugarcane planthopper *Eumetopina flavipes* (Anderson et al., 2013) as well as *Culicoides* biting midges (Eagles et al., 2014). Pests can move into the Torres Strait by natural processes (wind currents) as well as human activities (vessel movements, fishing activity) (Kompas et al., 2015).

## 5 CONCLUSIONS

In summary, our SNP-based analyses of *L. sativae* provides evidence of connections among populations that are mostly consistent with historical observations. Our study is the first to apply high-density SNPs markers to determine the population structure of *Liriomyza* and provide a baseline and foundation to further track leafminer movements across the region and further into the Australian mainland. The data provide a basis for quarantine detections to identify local versus international sources (c.f. Schmidt et al., 2019) and an ongoing assessment of new incursions can indicate risks associated with pesticide resistance genes entering local populations.

## ACKNOWLEDGEMENTS

We thank Ching-Chin Chien, Colby Maeda, Elia I. Pirtle, Glenn Bellis, I Wayan Supartha, Jessica Lye, John Trumble, Komivi Senyo Akutse, Kris Wyckhuys, Lihsin Wu, Mani Mua, Olivia Reynolds, Sally Cowan, Scott Ferguson, Shiuh-Feng Shiao and Wanxue Liu for generously sourcing international samples for this paper. In particular, we thank Mallik Malipatil for confirmation of morphological identifications. We thank LiJun Cao for bioinformatics assistance. This research was supported by RDE program for control, eradication and preparedness for vegetable leafminer (MT16004), funded by Hort Innovation.

## DATA AVAILABILITY STATEMENT

The data that support the findings of this study are available from the supplementary material, or the corresponding author upon reasonable request.

## REFERENCES

Alexander, D. H., & Lange, K. (2011). Enhancements to the ADMIXTURE algorithm for individual ancestry estimation. BMC Bioinformatics, 12(1), 1–6.

Alexander, D. H., Novembre, J., & Lange, K. (2009). Fast model-based estimation of ancestry in unrelated individuals. Genome Research, 19(9), 1655–1664.

Andersen, A., Nordhus, E., Thanq, V. T., An, T. T. T., Hung, H. Q., & Hofsvanq, T. (2002). Polyphagous *Liriomyza species* (Diptera: Agromyzidae) in vegetables in Vietnam. Tropical Agriculture, 79(4), 241–246.

Anderson, C. D., Epperson, B. K., Fortin, M. J., Holderegger, R., James, P. M., Rosenberg, M. S.,… & Spear, S. (2010). Considering spatial and temporal scale in landscape-genetic studies of gene flow. Molecular Ecology, 19(17), 3565–3575.

Anderson, K. L., & Congdon, B. C. (2013). Population genetics suggest that multiple invasion processes need to be addressed in the management plan of a plant disease vector. Evolutionary Applications, 6(4), 660–672.

Ballard, J. W. O., & Whitlock, M. C. (2004). The incomplete natural history of mitochondria. Molecular Ecology, 13(4), 729–744.

Beebe, N. W., Ambrose, L., Hill, L. A., Davis, J. B., Hapgood, G., Cooper, R. D.,… & van den Hurk, A. F. (2013). Tracing the tiger: population genetics provides valuable insights into the *Aedes (Stegomyia) albopictus* invasion of the Australasian Region. PLoS Neglected Tropical Diseases, 7(8), e2361.

Ben Abdelkrim, A., Hattab, T., Fakhfakh, H., Belkadhi, M. S., & Gorsane, F. (2017). A landscape genetic analysis of important agricultural pest species in Tunisia: The whitefly *Bemisia tabaci*. PLoS One, 12(10), e0185724.

Blacket, M. J., Rice, A. D., Semeraro, L., & Malipatil, M. B. (2015). DNA-based identifications reveal multiple introductions of the vegetable leafminer *Liriomyza sativae* (Diptera: Agromyzidae) into the Torres Strait Islands and Papua New Guinea. Bulletin of Entomological Research, 105(5), 533–544.

Blanchard, E. E. (1938). Descripciones y anotaciones de dípteros argentinos. Anales de la Sociedad Científica Argentina, 126(5), 345–386.

Broquet, T., Ray, N., Petit, E., Fryxell, J. M., & Burel, F. (2006). Genetic isolation by distance and landscape connectivity in the American marten (*Martes americana*). Landscape Ecology, 21(6), 877–889.

Carolina, Herr, J. C., & Johnson, M. W. (1992). Host plant preference of *Liriomyza sativae* (Diptera: Agromyzidae) populations infesting green onion in Hawaii. Environmental Entomology, 21(5), 1097–1102.

Catchen, J., Hohenlohe, P. A., Bassham, S., Amores, A., & Cresko, W. A. (2013). Stacks: an analysis tool set for population genomics. Molecular Ecology, 22(11), 3124–3140.

Catchen, J. M., Amores, A., Hohenlohe, P., Cresko, W., & Postlethwait, J. H. (2011). Stacks: building and genotyping loci de novo from short-read sequences. G3: Genes, Genomes, Genetics, 1(3), 171–182.

Chen, Y. & Zhao J. 1999. Influence of extreme high temperature on the biology of vegetable leafminer. Entomologia Sinica 6, 164–170.

Chunlin, W., Zongyi, Z., & Youling, H. (2005). Harmonization of national phytosanitary regulations in China. In Identification of risks and management of invasive alien species using the IPPC framework doi: http://www.fao.org/docrep/008/y5968e/y5968e0e.htm

Chen, M. Z., Cao, L. J., Li, B. Y., Chen, J. C., Gong, Y. J., Yang, Q.,… & Wei, S. J. (2021). Migration trajectories of the diamondback moth *Plutella xylostella* in China inferred from population genomic variation. Pest Management Science, 77(4), 1683–1693.

Cordeiro, E. M., Campbell, J. F., Phillips, T., & Akhunov, E. (2019). Isolation by distance, source-sink population dynamics and dispersal facilitation by trade routes: impact on population genetic structure of a stored grain pest. G3: Genes, Genomes, Genetics, 9(5), 1457–1468.

Coquilleau, M. P., Xu, X., Ridland, P. M., Umina, P. A., & Hoffmann, A. A. (2021). Variation in sex ratio of the leafminer *Phytomyza plantaginis* Goureau (Diptera: Agromyzidae) from Australia. Austral Entomology, 60(3), 610–620.

Dray, S., & Dufour, A. B. (2007). The ade4 package: implementing the duality diagram for ecologists. Journal of Statistical Software, 22(4), 1–20.

Eagles, D., Melville, L., Weir, R., Davis, S., Bellis, G., Zalucki, M. P.,… & Durr, P. A. (2014). Long-distance aerial dispersal modelling of *Culicoides* biting midges: case studies of incursions into Australia. BMC Veterinary Research, 10(1), 1–10.

Endersby-Harshman, N. M., Schmidt, T. L., Chung, J., van Rooyen, A., Weeks, A. R., & Hoffmann, A. A. (2020). Heterogeneous genetic invasions of three insecticide resistance mutations in Indo-Pacific populations of *Aedes aegypti* (L.). Molecular Ecology, 29(9), 1628–1641.

Fenoglio, M. S., Videl, M., Salvo, A., & Morales, J. M. (2019). Dispersal of the pea leaf miner *Liriomyza huidobrensis* (Blanchard, 1926) (Diptera: Agromyzidae): a field experiment. Revista de la Facultad de Ciencias Agrarias. Universidad Nacional de Cuyo, 51(2), 343–352.

Fola, A. A., Kattenberg, E., Razook, Z., Lautu-Gumal, D., Lee, S., Mehra, S.,… & Barry, A. E. (2020). SNP barcodes provide higher resolution than microsatellite markers to measure *Plasmodium vivax* population genetics. Malaria Journal, 19(1), 1–15.

Gitonga, Z. M., Chabi-Olaye, A., Mithöfer, D., Okello, J. J., & Ritho, C. N. (2010). Control of invasive *Liriomyza* leafminer species and compliance with food safety standards by small scale snow pea farmers in Kenya. Crop Protection, 29(12), 1472–1477.

Hardy, D. E., & Delfinado, M. D. (1980). Insects of Hawaii. In: Volume 13. Diptera: Cyclorrhapha III. Honolulu: University Press of Hawaii.

Hijmans, R. J., Williams, E., Vennes, C., & Hijmans, M. R. J. (2017). Package ‘geosphere’. Spherical Trigonometry, 1(7).

Hulme, P. E. (2009). Trade, transport and trouble: managing invasive species pathways in an era of globalization. Journal of Applied Ecology, 46(1), 10–18.

Hoffmann, A. A., White, V. L., Jasper, M., Yagui, H., Sinclair, S. J., & Kearney, M. R. (2021). An endangered flightless grasshopper with strong genetic structure maintains population genetic variation despite extensive habitat loss. Ecology and Evolution, 11(10), 5364–5380.

(IPPC) International Plant Protection Convention. (2017). Detection of *Liriomyza sativae* in Far North Queensland. IPPC Pest Report, AUS 80/1. Available from URL: https://www.ippc.int/en/countries/Australia/pestreports/2017/04/detection-of-liriomyza-sativae-in-far-north-queensland/ [Accessed 04 August 2021].

Jasper, M. E., Hoffmann, A. A., & Schmidt, T. L. (2021). Estimating dispersal using close kin dyads: The kindisperse R package. Molecular Ecology Resources. doi: 10.4049/jimmunol.1700729

Johnson, M. W. (2005). Our war with the insects: analysis of lost battles. In: K. M. Heinz, R. Frisbie & C. Borgan (Eds). Challenges within Entomology: A Celebration of the Past 100 years and a Look to the Next Century (pp. 207–223). College Station, TX: Texas A&M Press.

Johnson, M. W., Welter, S. C., Toscano, N. C., Ting, P., & Trumble, J. T. (1983). Reduction of tomato leaflet photosynthesis rates by mining activity of *Liriomyza sativae* (Diptera: Agromyzidae). Journal of Economic Entomology, 76(5), 1061–1063.

Jombart, T. (2008). adegenet: a R package for the multivariate analysis of genetic markers. Bioinformatics, 24(11), 1403–1405.

Kompas, T., Ha, P., & Spring, D. (2015). Baseline ‘consequence measures’ for Australia from the Torres Strait Islands pathway to Queensland: Papaya fruit fly, citrus canker and rabies. A Report Prepared for the Department of Agriculture and Water Resources (pp. 89). CEBRA, University of Melbourne and ACBEE, Australian National University.

Ma, D., Gorman, K., Devine, G., Luo, W., & Denholm, I. (2007). The biotype and insecticide-resistance status of whiteflies, *Bemisia tabaci* (Hemiptera: Aleyrodidae), invading cropping systems in Xinjiang Uygur Autonomous Region, northwestern China. Crop Protection, 26(4), 612–617.

Maino, J. L., Pirtle, E. I., Ridland, P. M., & Umina, P. A. (2019). Forecasting the potential distribution of the invasive vegetable leafminer using ‘top-down’and ‘bottom-up’models. bioRxiv doi:https://doi.org/10.1101/866996

Martinez, M. (1994). Un nouveau ravageur menace la région orientale: *Liriomyza sativae* Blanchard (Díptera, Agromyzidae). Bulletin de la Société Entomologique de France, 99(4), 356.

Martinez, M. & Bordat, D. (1996). Note sur la presence de *Liriomyza sativae* Blanchard au Soudan et an Cameroun (Diptera, Agromyzidae). Bulletin de la Société Entomologique de France, 101(1), 71–73.

Mason, G. A., Johnson, M. W., & Tabashnik, B. E. (1987). Susceptibility of *Liriomyza sativae* and *L. trifolii* (Diptera: Agromyzidae) to permethrin and fenvalerate. Journal of Economic Entomology, 80(6), 1262–1266.

Minkenberg, O. P. J. M. (1988). Dispersal of *Liriomyza trifolii*. EPPO Bulletin, 18(1), 173–182.

Murphy, S. T., & LaSalle, J. (1999). Balancing biological control strategies in the IPM of New World invasive *Liriomyza* leafminers in field vegetable crops. Biocontrol News and Information, 20, 91N–104N.

Musundire R, Chabi-Olaye A, Löhr B & Krüger K. 2011. Diversity of Agromyzidae and associated hymenopteran parasitoid species in the Afrotropical region: implications for biological control. BioControl 56, 1–9.

Oatman, E. R., & Kennedy, G. G. (1976). Methomyl induced outbreak of *Liriomyza sativae* on tomato. Journal of Economic Entomology, 69(5), 667–668.

Parcon, H., Loke, M., & Leung, P. (2010). Costs of transporting fresh fruits and vegetables to Honolulu from Hilo and Los Angeles. Honolulu (HI): University of Hawaii. 9 p. (Economic Issues EI-18).

Parish, J. B., Carvalho, G. A., Ramos, R. S., Queiroz, E. A., Picanço, M. C., Guedes, R. N., & Corrêa, A. S. (2017). Host range and genetic strains of leafminer flies (Diptera: Agromyzidae) in eastern Brazil reveal a new divergent clade of *Liriomyza sativae*. Agricultural and Forest Entomology, 19(3), 235–244.

Parrella, M. P. (1982). A review of the history and taxonomy of economically important serpentine leafminers *(Liriomyza* spp.) in California (Diptera: Agromyzidae). Pan-Pacific Entomologist, 58(4), 302–308.

Parrella, M. P. (1987). Biology of *Liriomyza*. Annual Review of Entomology, 32(1), 201–224.

Parrella, M. P., & Keil, C. B. (1984). Insect pest management: the lesson of *Liriomyza*. Bulletin of the Entomological Society of America, 30(2), 22–25.

Peterson, B. K., Weber, J. N., Kay, E. H., Fisher, H. S., & Hoekstra, H. E. (2012). Double digest RADseq: an inexpensive method for de novo SNP discovery and genotyping in model and non-model species. PloS One, 7(5), e37135.

Rašić, G., Filipović, I., Weeks, A. R., & Hoffmann, A. A. (2014). Genome-wide SNPs lead to strong signals of geographic structure and relatedness patterns in the major arbovirus vector, *Aedes aegypti*. BMC Genomics, 15(1), 1–12.

Reitz, S. R., Gao, Y., & Lei, Z. (2013). Insecticide use and the ecology of invasive *Liriomyza* leafminer management. In: S Trdan (Ed.), Insecticides - Development of Safer and More Effective Technologies (pp. 233–253). Rijeka: InTech. doi: 10.5772/53874

Rauf, A., Shepard, B. M., & Johnson, M. W. (2000). Leafminers in vegetables, ornamental plants and weeds in Indonesia: surveys of host crops, species composition and parasitoids. International Journal of Pest Management, 46(4), 257–266.

Ridland, P. M., Umina, P. A., Pirtle, E. I., & Hoffmann, A. A. (2020). Potential for biological control of the vegetable leafminer, *Liriomyza sativae* (Diptera: Agromyzidae), in Australia with parasitoid wasps. Austral Entomology, 59(1), 16–36.

Rousset, F., Lopez, J., & Belkhir, K. (2020). Package ‘genepop’. R package version, 1, 17.

Ryan, S. F., Lombaert, E., Espeset, A., Vila, R., Talavera, G., Dincă, V.,… & Shoemaker, D. (2019). Global invasion history of the agricultural pest butterfly *Pieris rapae* revealed with genomics and citizen science. Proceedings of the National Academy of Sciences, 116(40), 20015–20024.

Scheffer, S. J., & Lewis, M. L. (2005). Mitochondrial phylogeography of vegetable pest *Liriomyza sativae* (Diptera: Agromyzidae): divergent clades and invasive populations. Annals of the Entomological Society of America, 98(2), 181–186.

Spencer, K. A. (1973). Agromyzidae (Diptera) of Economic Importance. Springer Science & Business Media.

Schmidt, T. L., van Rooyen, A. R., Chung, J., Endersby-Harshman, N. M., Griffin, P. C., Sly, A.,… & Weeks, A. R. (2019). Tracking genetic invasions: Genome-wide single nucleotide polymorphisms reveal the source of pyrethroid-resistant *Aedes aegypti* (yellow fever mosquito) incursions at international ports. Evolutionary Applications, 12(6), 1136–1146.

Schmidt, T. L., Swan, T., Chung, J., Karl, S., Demok, S., Yang, Q.,… & Hoffmann, A. A. (2021). Spatial population genomics of a recent mosquito invasion. Molecular Ecology, 30(5), 1174–1189.

Tang, X. T., Ji, Y., Chang, Y. W., Shen, Y., Tian, Z. H., Gong, W. R., & Du, Y. Z. (2016). Population genetic structure and migration patterns of *Liriomyza sativae* in China: moderate subdivision and no Bridgehead effect revealed by microsatellites. Bulletin of Entomological Research, 106(1), 114–123.

Uchibori-Asano, M., Jouraku, A., Uchiyama, T., Yokoi, K., Akiduki, G., Suetsugu, Y.,… & Shinoda, T. (2019). Genome-wide identification of tebufenozide resistant genes in the smaller tea tortrix, *Adoxophyes honmai* (Lepidoptera: Tortricidae). Scientific Reports, 9(1), 1–12.

Waterhouse, D. F., & Norris, K. R. (1987). *Liriomyza* species (Diptera: Agromyzidae) leafminers. Biological control: Pacific prospects, 159–176.

White, C., Selkoe, K. A., Watson, J., Siegel, D. A., Zacherl, D. C., & Toonen, R. J. (2010). Ocean currents help explain population genetic structure. Proceedings of the Royal Society B: Biological Sciences, 277(1688), 1685–1694.

Xu, X., Coquilleau, M. P., Ridland, P. M., Umina, P. A., Yang, Q., & Hoffmann, A. A. (2021a). Molecular identification of leafmining flies from Australia including new *Liriomyza* outbreaks. Journal of Economic Entomology, 114(5), 1983–1990.

Xu, X., Ridland, P. M., Umina, P. A., Gill, A., Ross, P. A., Pirtle, E., & Hoffmann, A. A. (2021b). High incidence of related *Wolbachia* across unrelated leaf-mining Diptera. Insects, 12(9), 788.

Yamashita, H., Katai, H., Kawaguchi, L., Nagano, A. J., Nakamura, Y., Morita, A., & Ikka, T. (2019). Analyses of single nucleotide polymorphisms identified by ddRAD-seq reveal genetic structure of tea germplasm and Japanese landraces for tea breeding. PloS One, 14(8), e0220981.

Yan, J., Vétek, G., Pal, C., Zhang, J., Gmati, R., Fan, Q. H.,… & Li, D. (2021). ddRAD sequencing: an emerging technology added to the biosecurity toolbox for tracing the origin of brown marmorated stink bug, *Halyomorpha halys* (Hemiptera: Pentatomidae). BMC Genomics, 22(1), 1–15.

Yang, Q., Umina, P. A., Rašić, G., Bell, N., Fang, J., Lord, A., & Hoffmann, A. A. (2020). Origin of resistance to pyrethroids in the redlegged earth mite *(Halotydeus destructor)* in Australia: repeated local evolution and migration. Pest Management Science, 76(2), 509–519.

Zitter, T. A., & Tsai, J. H. (1977). Transmission of three potyviruses by the leafminer *Liriomyza sativae* (Diptera: Agromyzidae). Plant Disease Reporter, 61(12), 1025–1029.

