## Supplementary material for "Genome-wide SNPs of vegetable leafminer, *Liriomyza sativae*: insights into the recent Australian invasion": Xuefen_Supplementary Information.pdf

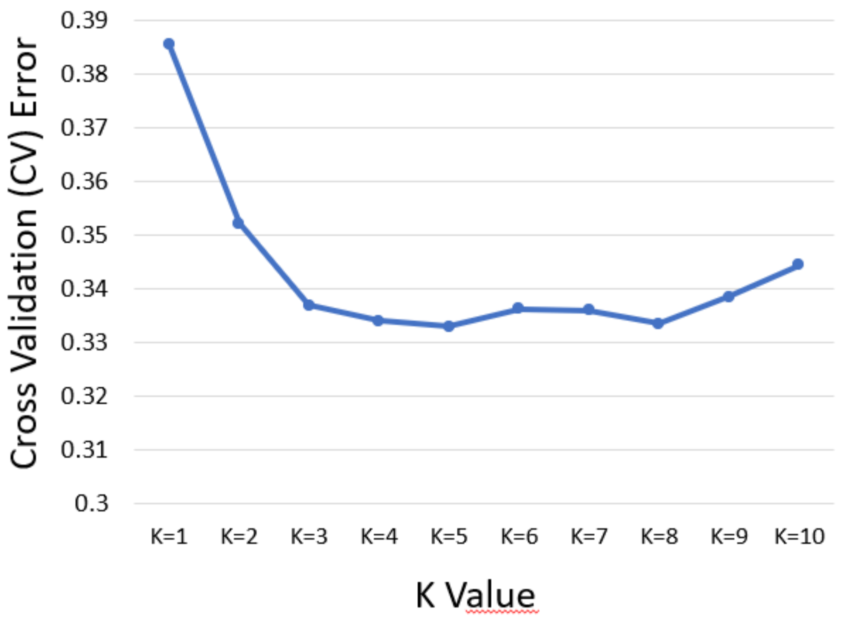


Fig. S1 Rate of change in cross-validation (CV) error between successive k-values (k-values ranged from 1 to 10, the minimum K is 5).

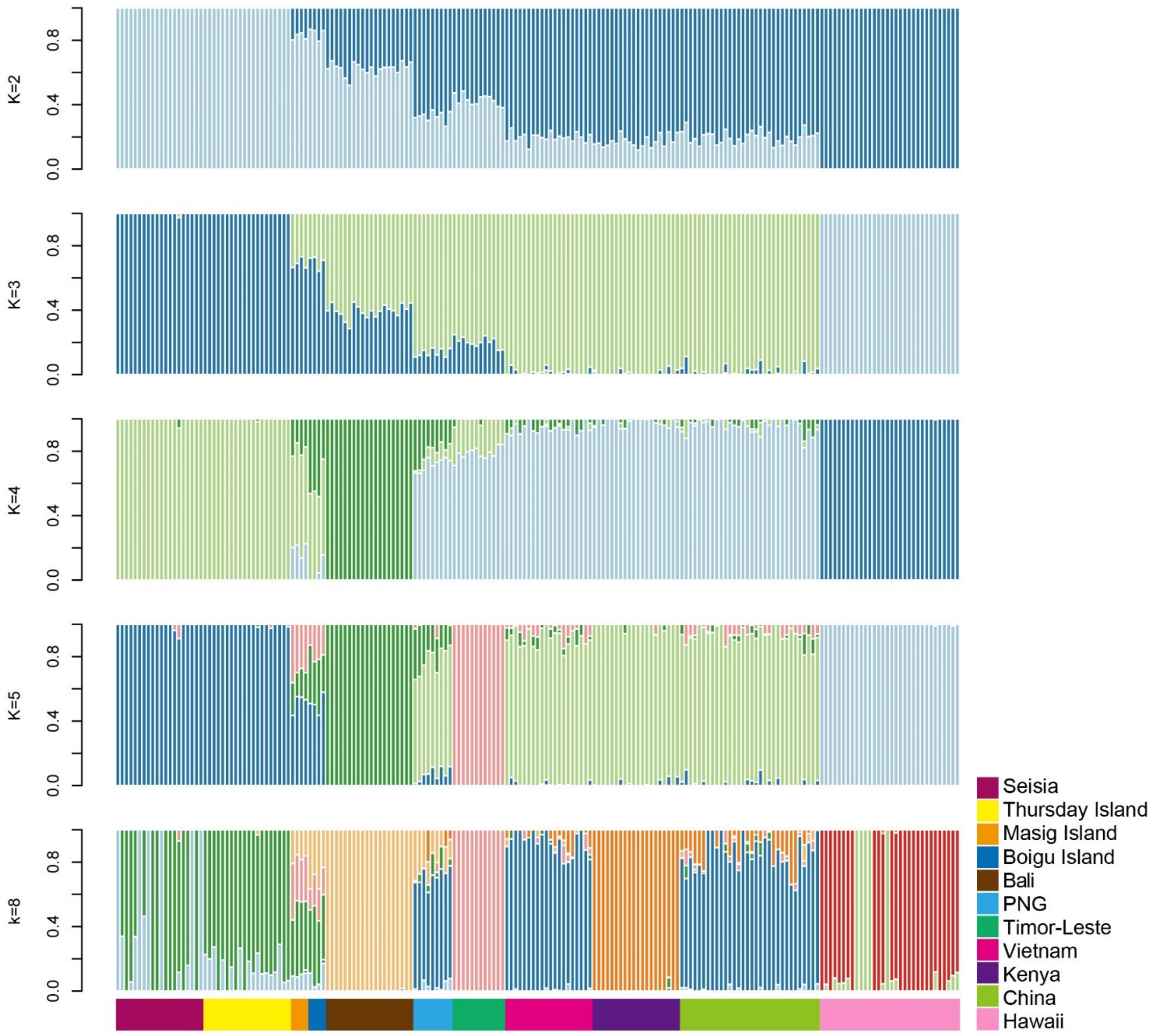


Fig. S2 Population structure of *L. sativae* based on 193 individuals inferred using Admixture analysis, the second minimum K is 8.

Table S1 and Table S2 see excel file.

Table S3 *Liriomyza sativae* in Hawaii – first recorded in 1921. Lonsdale (2011) lists the synonyms for *L. sativae*. In Hawaii, Frick (1952) described 4 new species of Agromyzidae. Three species (*L. canomarginis*, *L. minutiseta* and *L. pullata*) have subsequently been synonymised with *L. sativae*, while *L. hawaiiensis* is now synonymised with *L. brassicae* (Spencer 1973). Frick was not using male genitalia as a character and was using several inherently variable characters which have been shown to be not useful to delimit species (Hardy & Delfinado 1980).

| Species | Specimens used by Frick (1952) |
|---|---|
| *Liriomyza canomarginis* | Holotype ♂: Kaimuki, Oahu, **April 12, 1921**, O. H. Swezey, collector, ex *Indigofera* sp., deposited in the Hawaiian Sugar Planters' Association Experiment Station collection. Paratypes: 2♂♂, 5 ♀♀, Lualualei, Oahu,<br>June 26, 1943, T. Nishida, collector, ex bean; |
| *Liriomyza minutiseta* | Holotype ♂: Honolulu, Oahu, September 7, 1951, W. C. Mitchell, collector, ex tomato, deposited in the Hawaiian Sugar Planters' Association Experiment Station collection.<br>Paratypes: 2 ♂ ♂, 1 ♀, topotypical; 3 ♂ ♂,<br>Waianae, Oahu, January, 1951, W. C. Mitchell, collector, ex tomato;<br>3 ♂ ♂, 1 ♀, Waianae, Oahu, March, 1951, W. C. Mitchell, collector, ex tomato;<br>1 ♂, 2 ♀ ♀, Honolulu, Oahu, September 7, 1951, W. C. Mitchell, collector, ex eggplant;<br>1 ♂, 3♀ ♀, Honolulu, Oahu, September 7, 1951, W. C. Mitchell, collector, ex cauliflower;<br>2 ♂ ♂, Kunia, Oahu, September, 1951, D. E. Hardy, collector, ex squash. |
| *Liriomyza pullata* | Holotype ♀: Kanoa, Molokai, **March 3, 1929**, O. H. Swezey, collector, ex *Datura* sp., deposited in the Hawaiian Sugar Planters' Association Experiment Station collection.<br>Paratypes: 1 ♀, topotypical [collected from the type locality];<br>1 ♀, Makolelau, Molokai, March 23, 1929, O. H. Swezey, collector, ex *Lipochaeta* sp.;<br>1 ♀, Honolulu, Oahu, September, 1951, W. C. Mitchell, collector, ex *Aster* sp.;<br>1 ♂, 1 ♀, Waimanalo, Oahu, January 31, 1951, D. E. Hardy, collector, sweeping;<br>1 ♀, Honolulu, Oahu, April, 1951, D. E. Hardy, collector, at light. |

### References

Frick, K. E. (1952). Four new Hawaiian *Liriomyza* species and notes on other Hawaiian Agromyzidae (Diptera). *Proceedings of the Hawaiian Entomological Society* 14, 509–518.

Hardy DE & Delfinado MD. 1980. Family Agromyzidae In *Insects of Hawaii*. *Volume 13, Diptera: Cyclorrhapha III, Series Schizophora Section Acalypterae, Exclusive of Family Drosophilidae* (eds DE Hardy & MD Delfinado). pp. 190-222. The University Press of Hawaii, Honolulu.

Lonsdale O. 2011. The *Liriomyza* (Agromyzidae: Schizophora: Diptera) of California. *Zootaxa* 2850, 1–123.

Spencer KA. 1973. Agromyzidae (Diptera) of Economic Importance. In: E. Schimitschek (Ed.), *Series Entomologica*, Vol. 9, Springer-Science+Business Media, Dordrecht, The Netherlands.
